# EEG Microstates Reveal Differential Network Dynamics under Constant Current and Oscillatory Brain Stimulation

**DOI:** 10.1101/2025.10.08.681119

**Authors:** Inga Griškova-Bulanova, Povilas Tarailis, Marko Živanović, Saša Filipović, Jovana Bjekić

## Abstract

**Background:** EEG microstates are brief, quasi-stable scalp topographies that index large-scale network dynamics and may sensitively capture the net effects of transcranial electrical stimulation (tES).

**Objective:** To evaluate differential effects of three types of tES - tDCS, tACS and oscillatory tDCS - (otDCS), on canonical EEG microstates (A-D) in healthy adults.

**Methods:** In a randomized, sham-controlled, crossover study, 42 participants completed four sessions (tDCS, tACS, otDCS, sham). Stimulation (20 min) used a P3–cheek montage: tDCS +1.5 mA; tACS at individualized theta frequency (ITF, 4–8 Hz), ±1 mA; otDCS anodal with ±0.5 mA oscillation around +1.5 mA at ITF. A five-minute resting EEG (eyes closed then eyes open) was recorded pre- and post-intervention. Microstates were extracted (A–D), back-fitted, and assessed on duration, occurrence, contribution, and mean GFP using linear mixed-effects models with sham and pre/post adjustments.

**Results:** Four canonical microstates explained ~80% variance with stable topographies across conditions. Modulation patterns were modality-specific. MS A (sensory/arousal) increased across all active protocols, strongest after otDCS. MS B (visual-autobiographical) was consistently suppressed, again most following otDCS. MS C (self-referential) decreased selectively after oscillatory stimulation (tACS, otDCS) only. MS D (executive/attention) diverged by waveform: enhanced by tACS and otDCS but reduced by tDCS. Across outcomes, otDCS produced the largest and most widespread effects, overlapping features of both tDCS (tonic/stabilizing) and tACS (oscillatory/entraining) influences.

**Conclusions:** Resting-state EEG microstates provide a sensitive systems-level assay of tES aftereffects. Constant and oscillatory current waveforms reorganize network states in dissociable ways, with otDCS exerting the most robust, comprehensive modulation. These findings support microstates as practical biomarkers for differentiating, optimizing, and monitoring neuromodulation strategies.

## Introduction

Low-intensity transcranial electrical stimulation (tES) has become widely used for investigating and modulating human brain function in a non-invasive manner. By delivering weak currents through the scalp, tES protocols can interact with ongoing neuronal activity in distinct ways. Transcranial direct current stimulation (tDCS) induces polarity-dependent shifts in cortical excitability via constant currents (1–3), while transcranial alternating current stimulation (tACS) applies rhythmic stimulation to entrain, shift, or interfere with endogenous oscillations (4–6). Oscillatory transcranial direct current stimulation (otDCS) combines these approaches by embedding a sinusoidal component within a steady DC offset, potentially integrating excitability shifts with oscillatory entrainment (7–9).

Effective use of tES depends on the precise understanding of how externally applied currents reshape intrinsic patterns of brain activity. A central challenge lies in uncovering how different stimulation modalities affect large-scale brain dynamics. Resting state EEG data allow for identification of microstates, that is, brief, quasi-stable topographies of brain activity lasting tens to hundreds of milliseconds (10). Microstates are typically classified into four canonical classes (A, B, C, and D), each characterized by a distinct scalp topography and functional association. Their temporal parameters, including duration, occurrence, and contribution, are increasingly recognized as indices of functional brain states linked to cognition and clinical conditions (10).

EEG microstates offer a powerful framework for tracking the effects of brain stimulation because they provide a temporally precise index of large-scale brain network dynamics (10,11). Unlike traditional oscillatory analyses that focus on local frequency-specific activity, microstates capture transient, quasi-stable topographies that reflect the coordinated activity of distributed functional networks such as the salience, default mode, attention, and executive control systems (12,13). This network-level sensitivity makes microstates particularly suitable for neuromodulation studies, where interventions such as tDCS, tACS, otDCS, repetitive transcranial magnetic stimulation (rTMS), or intermittent theta-burst stimulation (iTBS) are expected to induce network-level reorganizations rather than isolated local changes (14–16). Across both clinical and healthy populations, microstate parameters have been shown to reliably differentiate between pathological and normal brain function (17) and to normalize following effective stimulation, closely paralleling improvements in pain, mood, cognition, or psychiatric symptoms (18–20). Their sensitivity to both stimulation-induced reorganization and clinical outcome highlights microstates as cost-effective, accessible biomarkers that can bridge mechanistic understanding and therapeutic monitoring of brain stimulation effects.

Despite this potential and their growing application in clinical neuroscience, evidence on how microstates reflect the effects of brain stimulation remains scarce. Initial findings with tDCS suggest that microstates are sensitive to stimulation-induced changes: in healthy participants, tDCS over motor and prefrontal regions reorganized network states linked to pain and anxiety (21), while in autism spectrum disorder and depression, stimulation over the DLPFC increased the presence of functionally relevant microstates and aligned with symptom improvement (18,22). In obsessive-compulsive disorder, orbitofrontal tDCS normalized aberrant microstate transitions, with microstate C emerging as a potential biomarker of therapeutic response (23). More recently, in schizophrenia, repeated HD-tDCS delivered during working memory training increased the presence of MS B, which was reduced at baseline relative to controls, indicating a normalization of disorder-specific network alterations (24). These early studies highlight microstates as a promising, yet still underexplored, approach to track the large-scale brain effects of tDCS.

Even fewer studies have examined the effects of tACS on EEG microstates, but the initial findings indicate that this approach can reveal stimulation-induced network reorganization. In healthy adults, γ-tACS at 40 Hz over DLPFC was shown to reduce the presence of microstate C and increase microstates D and B, with these changes correlating with enhanced working memory performance (16). Similarly, another study reported a consistent decrease in microstate C and an increase in microstate D following γ-tACS, highlighting the sensitivity of microstate analysis to capture large-scale effects of rhythmic brain stimulation (25).

Together, these findings position EEG microstates as a promising yet underutilized tool for probing stimulation-induced changes in large-scale brain networks. To advance this line of research, we conducted the first within-subject, systematic comparison of constant and oscillatory tES modalities (tDCS, tACS, and otDCS) against sham, examining their effects on canonical EEG microstates. The study aims to identify modality-specific signatures of network reorganization, thereby providing novel insights into the network-level mechanisms of various tES modalities, while also evaluating the sensitivity of EEG microstates as a marker for differentiating the large-scale effects of distinct stimulation protocols.

## Methods

### Design

This study employed a sham-controlled, randomized, crossover tES-EEG experimental design and was conducted as part of the larger MEMORYST project (for details, see (26)). Namely, each participant completed four experimental sessions, at least one week apart to avoid potential carryover effects. In a counterbalanced order, they received one of four tES conditions: tDCS, otDCS, tACS, or sham. Stimulation was delivered concurrently with cognitive tasks aimed at keeping participants at a similar level and the type of cognitive engagement. Resting-state EEG was recorded for 5 minutes immediately before and after each stimulation, beginning with 3 minutes of eyes-closed, followed by 2 minutes of eyes-open recording. The study was conducted in accordance with the Declaration of Helsinki and approved by the Institutional Ethics Board (EO129/2020).

### Participants

The study recruited healthy young adults of both sexes who met the inclusion criteria and agreed to participate in exchange for monetary compensation (€8/hour). Individuals who expressed interest were first screened using an online eligibility self-assessment checklist. Inclusion criteria required participants to be right-handed, aged between 20 and 35 years, with normal or corrected-to-normal vision. Exclusion criteria included a history of seizures, neurological or psychiatric disorders, cognitive or learning disabilities, traumatic brain injury, metal implants in the head, chronic or acute dermatological conditions on the head, confirmed or suspected pregnancy, and current use of psychoactive substances or medications.

A total of 42 participants (26 female), aged 22–34 years (*M* = 25.05, *SD* = 3.55), were enrolled in the study. An a priori power analysis was conducted using G*Power software to determine the appropriate sample size. Assuming a within-subjects design with four conditions, α = .05, and power (1 − β) = 0.80, a minimum of 36 participants was required to detect a medium effect size (f = 0.25). The final sample size of 42 reflects deliberate oversampling to ensure adequate power despite potential attrition and signal quality issues common in tES-EEG research.

### Transcranial electrical stimulation and EEG acquisition

Starstim32 system (Neuroelectrics Inc., Barcelona, Spain), a lightweight, mobile, battery-powered device remotely operated via the Neuroelectrics® Instrument Controller (NIC2) software, was used for both tES delivery and EEG recording. The electrodes were secured using a neoprene cap with a 39-channel grid based on a subset of the international 10–10 EEG system. The EEG-tES setup is presented in Figure 1 (panel A), alongside the visualization of the induced e-fields (panel C).

**Figure 1.**
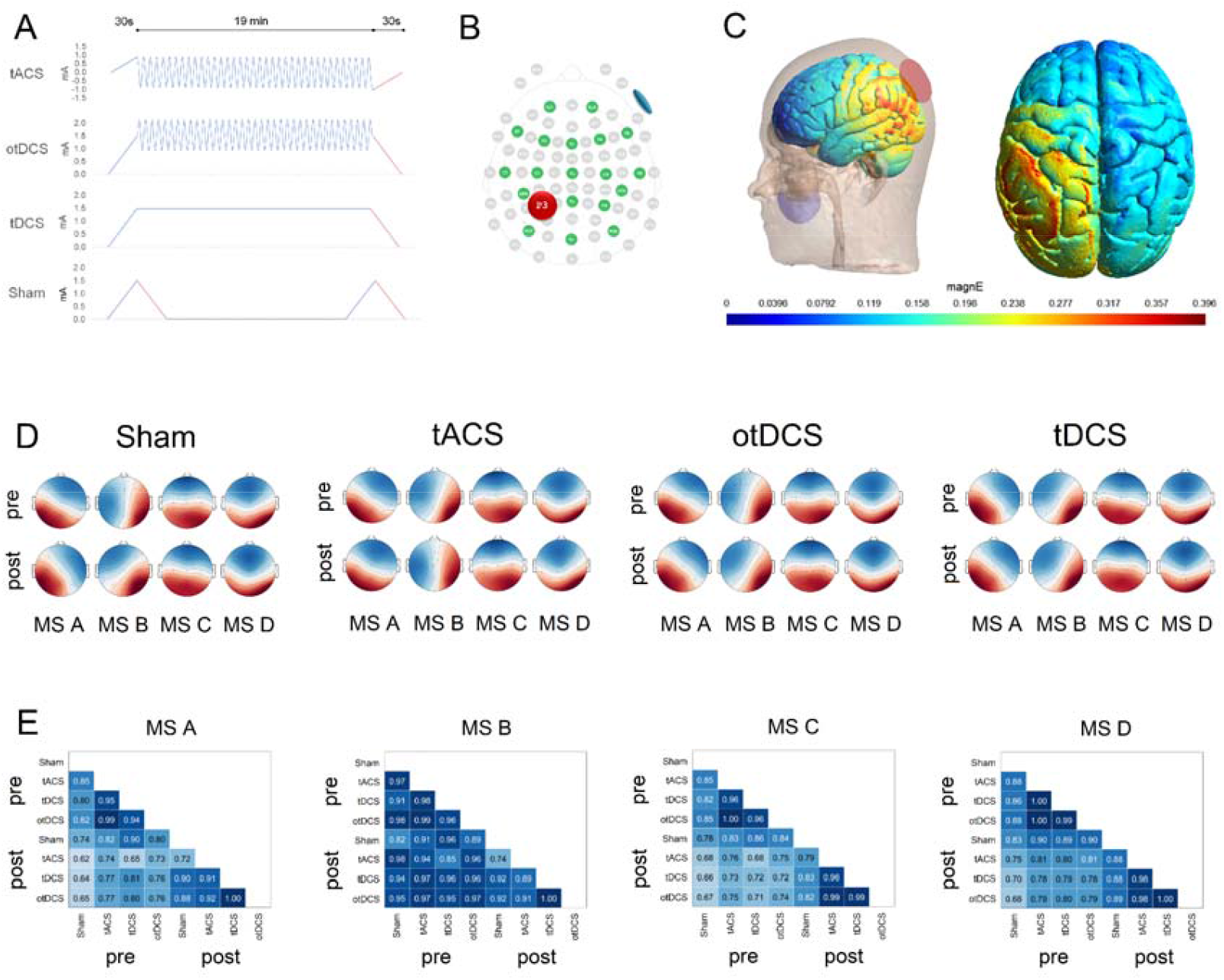
(A) Overview of stimulation protocols: tACS (30 s ramp up to 1 mA, 19 min at ITF, 30 s ramp down), otDCS (30 s ramp up to 1.5 mA, 19 min at 1.5 ± 0.5 mA at ITF, 30 s ramp down), tDCS (30 s ramp up to 1.5 mA, 19 min at 1.5 mA, 30 s ramp down), and sham (30 s ramp up/down to 1.5 mA, 19 min at 0 mA, followed by a second 30 s ramp up/down), (B) tES-EEG electrode setup and (C) SimNIBS simulation of induced electric field for 1.5mA current intensity. (D) The four microstate maps were extracted pre- and post-each stimulation condition: sham, tDCS, tACS, and otDCS. (C) Spatial correlation coefficients per microstate class between group-level topographies for all stimulation conditions

For stimulation, round rubber electrodes (25 cm^2^) were inserted into saline-soaked sponge pockets. One electrode was placed over the left posterior parietal cortex (P3), and the other electrode was fixed to the contralateral cheek using medical adhesive tape (Figure 1, panel B). Stimulation lasted 20 minutes across all conditions. Anodal tDCS was delivered as a constant +1.5 mA current. tACS was applied at the individual theta frequency (ITF, 4–8 Hz) as a ±1 mA sinusoidal waveform (2 mA peak-to-peak). otDCS used an anodal waveform oscillating ±0.5 mA around +1.5 mA (i.e., +1 to +2 mA) at ITF. All active protocols included 30-second ramp-up and ramp-down periods. In the sham condition, current was applied for 60 seconds at the beginning and end, using a 30-second ramp-up to 1.5 mA followed by a 30-second ramp-down.

EEG was recorded using 19 Ag/AgCl gel electrodes (4 mm diameter, 1 cm^2^ gel-contact area) positioned at Fp1, Fp2, Fz, F3, F4, F7, F8, T7, T8, Cz, C3, C4, CP5, CP6, Pz, P4, PO7, PO8, and Oz. This electrode configuration was chosen to provide broad cortical coverage while avoiding physical overlap with stimulation sites. Reference and ground were provided by either a dual CMS/DRL ear-clip electrode attached to the right earlobe or two pre-gelled sticktrodes placed on the right mastoid (CMS) and just below it (DRL). EEG signals were recorded at a sampling rate of 500 Hz, with a bandwidth of 0–125 Hz (DC-coupled), 24-bit resolution, and a voltage resolution of 0.05 µV.

### EEG analysis

The microstate analysis was performed using a microstate plugin for EEGLAB (27) and custom-written functions. For each condition (Sham, tDCS, tACS, otDCS), data were analyzed in two steps: at the individual and at the group levels. First, for each subject, Global Field Power (GFP) was calculated. To ensure a sufficient signal-to-noise ratio, only the maps at momentary peaks of the GFP were extracted (28,29) and submitted to a modified k-means clustering algorithm (30), ignoring polarity differences. For each subject, the number of clusters ranged between 2 and 8, and to maximize global explained variance (GEV), clusterization was repeated 50 times for each number of clusters. For the second step, individual topographies at each number of clusters were averaged across participant group-wise, using a permutation algorithm that maximizes the common variance between subjects (31). The optimal number of group-level topographies for each condition was estimated as 4 based on the Silhouette method (32,33). To quantify the spatial similarity between the groups, spatial correlation was calculated, again ignoring polarity differences.

Group-level topographies were backfitted to individual EEGs by a winner-takes-all approach: for each GFP peak, the scalp potential map was compared to each group-level topography and was assigned to the best-matching class based on the spatial correlation. After assigning a microstate class to each GFP peak, these labels were extended to all intervening time frames. The segment boundaries were defined at the midpoints between consecutive GFP peaks (closest neighbor approach), resulting in a continuous segmentation of the EEG.

Then, the following parameters were calculated for each microstate: Mean duration reflecting intracortical generators’ synchronous activity for each occurrence of a particular microstate configuration, and is measured in milliseconds (ms) (34); Occurrence rate referring to the mean number of times a microstate occurred during one second period and interpreted as the tendency of intracortical sources to be synchronously activated (measured in Hertz (Hz), (34); Coverage reflecting the total percent of the time frames for which a microstate is accounted (34,35); Global Explained Variance (GEV) standing for the sum of the explained variances weighted by the GFP at each moment in time and is measured in percentages (Murray et al., 2008).

### Statistical analysis

The effects of different tES modalities on EEG microstate dynamics were examined using a series of linear mixed-effects models (LMMs). In all models, stimulation CONDITION (real tES vs. sham) and TIME (pre vs. post) were entered as fixed factors, while participant ID was included as a random grouping factor (e.g., *Duration_MS1 ~ condition * time + (1* | *ID)*). Separate models were specified for each real tES–sham comparison, for each microstate class (A, B, C, D), and for each outcome variable (duration, occurrence, contribution, GFP). The primary focus was on the interaction terms, as these indicate stimulation-specific changes (i.e., whether pre-to-post differences varied between real and sham stimulation). All post hoc pairwise comparisons used Satterthwaite’s method for degrees of freedom estimation and were corrected for multiple testing with the Holm procedure.

To directly compare active tES modalities (tDCS, tACS, otDCS), outcomes were baseline-(e.g., *tACS_diff = tACS_post – tACS_pre*) and sham-adjusted (*tACS_diff – Sham_diff*) for each participant. These adjusted outcomes were then analyzed using LMMs with stimulation type (tDCS, tACS, otDCS) as a fixed factor and participant ID as a random factor. Post hoc comparisons were conducted with Holm correction to control for multiple comparisons. All statistical analyses were carried out using JASP (v0.95) and the lme4 package in R (v4.5.1).

## Results

### Microstate topographies and temporal parameters

Four canonical microstate classes - A, B, C, and D were identified, consistently explaining IZ80% of variance (range: 79.99% - 81.28%; without differences between tES conditions nor pre-to-post, all *p* > .136). Figure 1 shows the corresponding topographical representations of microstates in pre- and post-stimulation conditions (panel D). The spatial correlations between microstates extracted pre and post tES ranged between .72 and .97 in corresponding tES conditions and .65 to 1 for cross-stimulation (panel E). The descriptive characteristics of microstates, such as duration, occurrence, contribution, and GFP, are provided in the Supplementary Material (Table S1).

The full account of the 2 (CONDITION: real, sham) x 2 (TIME: pre, post) LMMs on EEG microstate measures (duration, occurrence, contribution, and GFP) is provided in the Supplementary material for each of the tES types (Table S2).

### tDCS effects

A significant CONDITION x TIME interaction was observed for MS A occurrence [*F*_*(1,123*)_ = 4.05, *p =* .*046*, η_p_^2^ = .03], and contribution [*F*_*(1,123*)_ = 6.07, *p =* .*015*, η_p_^2^ = .05] (see Figure S1 of the Supplementary material). At baseline, contribution was higher in the sham condition [*t(123) = 4*.*3*2, *p <*.001], with a similar trend observed for occurrence [*t*(123) = 2.23, *p =* .*055*]. Post stimulation, increases in MS A measures following tDCS relative to sham abolished the baseline differences (*p*-values > .402).

For MS B, significant interactions were found for duration [*F*_(1,123)_ = 24.53, *p* <.001, η_P_^2^ = .17], contribution [*F*_(1,123)_ = 4.18, *p* = .043, η_P_^2^ = .03], and GFP [*F*_(1,123)_ = 14.62, *p* <.001, η_P_^2^ = .11], Although baseline duration and contribution were higher in the tDCS condition (*p*-values <.002), the main post-tDCS effect was a marked reduction in duration [*t*(123) = 3.07, *p* = .003], relative to sham. Furthermore, with equal baselines, GFP was found to be lower post-tDCS relative to sham [*t*(123) = 4.05, *p* <.001].

For MS C, an interaction was observed for occurrence [*F*_(1,123)_ = 5.84, *p* = .017, η_P_^2^ = .05], and GFP [*F*_(1,123)_ = 6.48, *p* = .012, η_P_^2^ = .05]. With equal baselines (*p*-values > .371), higher occurrence [*t*(123) = 3.34, *p* =.002] and lower GFP [*t*(123) = 2.70, *p* = .016] were recorded following tDCS relative to sham.

Finally, for MS4 significant interaction was observed for duration [*F*_(1,123)_ = 17.11, *p* <.001, η_P_^2^ = .12], and GFP [*F*_[1.123)_ = 31.25, *p* <.001, η_P_^2^ = .20]. With equal baseline (*p-*values >.675), post-stimulation tDCS effects were consistently negative, with shorter duration [*t*(123) = 2.84, *p* = .011] and lower GFP [*t*(123) = 3.15, *p* = .004] relative to sham.

### tACS effects

For MS A, significant CONDITION x TIME interactions were observed for duration [*F*_(p1,123)_ = 5.77, *p* = .018, η_P_^2^ = .04], contribution [*F*_(1,123)_ = 5.86, p =.017, η_P_^2^ = .05], and GFP [*F*_(1,123)_ = 7.03, *p* = .010, η_P_^2^ = .05] (see Figure S2 of the Supplementary material). At baseline, no differences were found between tACS and sham (all *p-*values > .108). In contrast, post-stimulation tACS produced a significantly higher contribution [*t*(123) = 2.95, *p* = .008] and a trend toward longer duration [*t*(123) = 2.26, *p* = .051] compared with sham.

For MS B, significant interactions were found for duration [*F*_(1,123)_ =25.00, *p* <.001, η_P_^2^ =.17], contribution [*F*_(1,123)_ = 19.94, *p* <.001, η_P_^2^ = .14], and GFP [*F*_(1,123)_ = 5.01, *p* = .027, η_P_^2^ = .04]. For contribution [*t*(123) = 2.11, *p* = .037] and, on a trend-level duration [*t*(123) = 1.93, *p* = .056], unequal baselines were observed with higher pre-tACS than pre-sham values. In post-stimulation assessment, however, significantly lower values were observed for all three measures compared to sham [duration *t*(123) = 5.14, *p* <.001; contribution *t*(123) = 4.21, *p* <.001; GFP *t*(123) = 2.75, *p* =.014].

For MS C, significant interactions were found for the measures of duration [*F*_(1,123)_ = 6.52, *p* =.012, η_P_^2^ = .05] and contribution [*F*_(1,123)_ = 4.48, *p* = .036, η_P_^2^ = .04]. At the baseline, no differences between tACS and sham were observed (all *p*-values > .445), while both measures showed significantly lower post-stimulation values relative to sham [duration *t*(123) = 3.85, *p* <.001; contribution *t*(123) = 3.76, *p* <.001].

For MS D, significant interactions were found for occurrence [*F*_(1,123)_ = 6.04, *p* = .015, η_P_^2^= .05] and contribution [*F*_(1,123)_ = 4.70, *p* < .032, η_P_^2^ = .04]. At the baseline, no differences between tACS and sham were observed (all *p-*values > .847). Both measures showed higher post-simulation values relative to sham [occurrence *t*(123) = 3.28, *p* = .003; contribution *t*(123) = 3.10, *p* = .005].

### otDCS effects

For MS A significant CONDITION x TIME interactions were observed for measures of duration [*F*_(1,123)_ = 17.11, *p* <.001, η_P_^2^ = .12], occurrence [*F*_(1,123)_ = 16.79, *p* <.001, η_P_^2^ =.12], and contribution [*F*_(1,123)_ = 31.25, *p* <.001, η_P_^2^ = 20] (see Figure S3 of the Supplementary material). For all three measures, unequal baselines (all *p-*values <.003) with higher pre-sham than pre-otDCS values were observed. Conversely, after the stimulation, a higher occurrence [*t*(123) = 2.58, *p* = .011] was observed relative to sham, as well as a trend-level increase in contribution [*t*(123) = 1.77, *p* = .079].

For MS B, significant interactions were found for duration [*F*_(1,123)_ = 49.04, *p* <.001, η_P_^2^ =.29], contribution [*F*_(1,123)_ = 28.53, *p* <.001, η_P_^2^ =.19], and GFP [*F*_(1,123)_ = 19.87, *p* <.001, η_P_^2^ = .14]. For all three measures, unequal baselines (all *p*-values <.022) with higher pre-otDCS in comparison to pre-sham values were recorded. However, the inverse pattern was observed in post-assessment. Namely, shorter duration [*t*(123) = 3.15, *p* = .002] and lower GFP [*t*(123) = 3.98, *p* <.001] were observed following otDCS compared to sham.

For MS C, significant interactions were found for duration [*F*_(1,123)_ = 15.26, *p* <.001, η_P_^2^ =.11], contribution [*F*_(1,123)_ = 12.08, *p* <.001, η_P_^2^ = .09], and GFP [*F*_(1,123)_ = 5.62, *p* = .019, η_P_^2^ = .04]. At baseline, no differences were observed between otDCS and sham. Relative to sham, following otDCS, shorter duration [duration *t*(123) = 5.53, *p* <.001], as well as lower contribution [*t*(123) = 4.80, *p* <.001], and lower GFP [*t*(123) = 4.53, *p* <.001] were observed.

For MS D, significant interactions were found for occurrence [*F*_(1,123)_ = 16.54, *p* <.001, η_P_^2^ = .12] and contribution [*F*_(1,123)_ = 16.52, *p* <.001, η_P_^2^ = .12]. For both measures, no differences were observed between otDCS and sham at baseline (*p*-values > .646). At post-assessment, both occurrence [*t*(123) = 5.29, *p* <.001] and contribution [*t*(123) = 5.06, *p* <.001] showed an increase compared to sham.

### Comparison between tDCS, tACS, and otDCS effects

Across all tES modalities, MS A tended to increase its presence, but the magnitude of facilitation differed significantly between stimulation types (Figure 2). tACS produced greater facilitation than tDCS on GFP (*t*(82) = 3.19, *p* = .006), whereas otDCS outperformed tACS on both occurrence (*t*(82) = 3.76, *p* <.001) and contribution (*t*(82) = 3.38, *p* = .002). Compared with tDCS, otDCS showed stronger facilitatory effects on all MS A measures (duration: *t*(82) = 2.77, *p* = .021; occurrence: *t*(82) = 2.46, *p* = .032; contribution: (*t*(82) = 3.83, *p* <.001), and GFP: *t*(82) = 2.33, *p* = .045).

**Figure 2.**
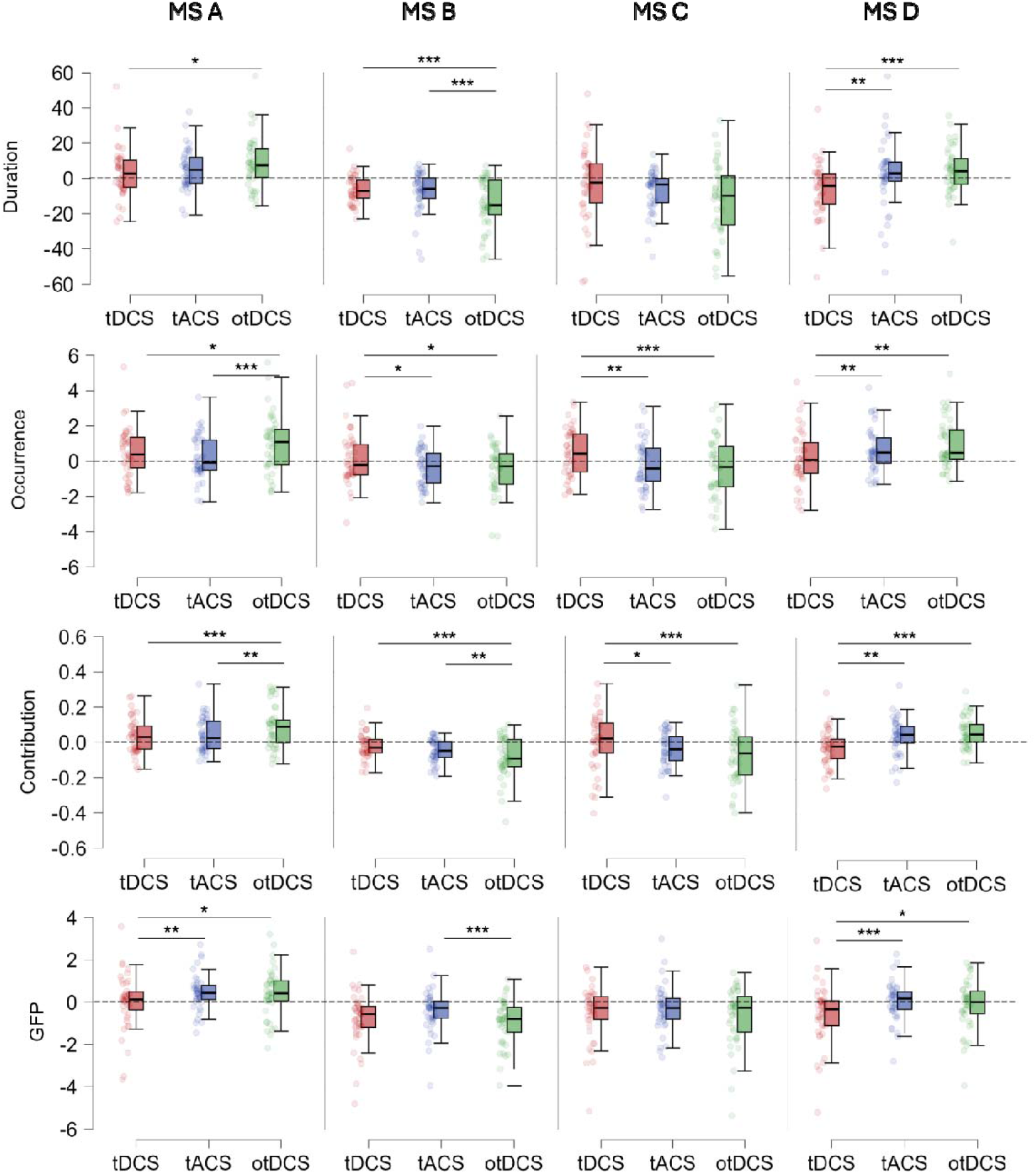
Differential effects of tDCS, tACS, and otDCS on EEG microstate parameters. \**p <*.*05*, \*\**p <*.*01*, \*\*\**p <*.*00*1. The dotted line represents a reference point for deviations from baseline-corrected sham.

The overall negative impact on MS B also varied by stimulation type. Relative to tDCS, tACS was associated with a lower occurrence (*t*(82) = 2.45, *p* = .033), while otDCS produced shorter duration (*t*(82) = 4.24, *p* <.001), lower occurrence (*t*(82) = 2.88, *p* = .015) lower contribution (*t*(82) = 4.72, *p* <.001). Compared with tACS, otDCS again exerted stronger effects, further decreasing duration (*t*(82) = 3.66, *p* <.001), contribution (*t*(82) = 2.85, *p* = .011), and GFP (t(82)=4.02, p<.001)

For MS C, the effects of both tACS and otDCS differed significantly from tDCS, each resulting in lowered occurrence (*t*(82) = 3.11, *p* = .005, *t*(82) = 3.99, *p* <.001, respectively) and contribution (*t*(82) = 2.30, *p* = .048, *t*(82) = 3.85, *p* <.001, respectively).

Pairwise comparisons confirmed the initially divergent effects of tES modalities on MS D. Namely, statistically significant differences between tDCS and tACS, as well as otDCS, were observed. tACS produced significantly greater duration (*t*(82) = 3.36, *p* = .002), occurrence (*t*(111.73) = 2.87, *p* = .010), contribution (*t*(82) = 3.23, *p* = .004), and GFP (*t*(82) = 3.96, *p* <.001) compared to tDCS. Finally, otDCS showed the largest facilitation overall, significantly exceeding tDCS on all measures (duration: *t*(82) = 4.18, *p* <.001), occurrence: *t*(82) = 3.02, *p* = .010; contribution: t(82) = 4.44, p <.001; and GFP: t(82) = 2.51, p = .028)), but differences between otDCS and tACS did not reach statistical significance.

## Discussion

EEG microstates provide a valuable framework for examining the net effects of brain stimulation, as they capture transient, quasi-stable topographies that reflect the coordinated activity of large-scale functional networks (13). This network-level perspective is particularly relevant in the context of tES, where low-intensity currents are hypothesized to induce widespread changes in cortical dynamics rather than isolated local effects, given the reduced focality of the applied electric fields (36–38). Although microstate analysis has been increasingly applied in psychiatric and neurological populations, few studies to date have explored how microstates respond to brain stimulation. The present study contributes to this emerging field by directly comparing the immediate aftereffects of the three distinct modalities of tES (tDCS, tACS, and otDCS) on the resting-state EEG microstates in healthy adults. We show that canonical microstates are sensitive to tES-induced neurophysiological changes, and that they are differentially modulated depending on stimulation modality, with otDCS consistently producing the largest effect sizes and overlapping with the effects of both tDCS and tACS. These findings highlight the sensitivity of EEG microstates to detect large-scale network reorganizations induced by brain stimulation and position otDCS as a particularly potent protocol for modulating global brain dynamics.

Across stimulation modalities (tDCS/tACS/otDCS), several consistent patterns emerged. MS A was facilitated by all active protocols, with otDCS producing the strongest increases across duration, occurrence, contribution, and GFP. MS B was consistently suppressed, again most strongly by otDCS, which reduced all parameters beyond the effects of tDCS or tACS. MS C was selectively reduced by theta-oscillatory stimulation (tACS and otDCS). Finally, MS D showed divergent effects: it was suppressed by tDCS but enhanced under theta-oscillatory protocols, with otDCS producing the strongest facilitation. Together, these findings indicate both shared and setting-specific influences of tES on microstate dynamics, with otDCS emerging as the most effective in modulating network-level brain activity.

Interpreting our results in functional terms provides further insight into how different stimulation settings shape brain dynamics. MS A, associated with auditory–visual processing and arousal (12,13), was facilitated across all protocols, suggesting that tES broadly enhances sensory readiness and alertness, which may translate into heightened vigilance to external input. This falls in line and explains findings of behavioral studies showing that parietal tDCS and tACS affect attention and vigilance (e.g., (39–41). MS B, linked to visual self-processing and autobiographical memory (11,13), was consistently suppressed, particularly under oscillatory protocols, indicating a shift away from internally oriented visual–autobiographical states toward externally focused processing. MS C, tied to self-referential and personally significant internal mentation (13,42), was selectively reduced by tACS and otDCS, reflecting a dampening of internally focused cognition and a reallocation of resources toward task-ready states. By contrast, MS D, associated with different cognitive processes - notably executive control and attention-shifting (13,43), was enhanced by tACS and otDCS but suppressed by tDCS, suggesting that theta-oscillatory protocols preferentially promote control-related engagement, whereas constant currents exert more stabilizing or inhibitory influences. According to Greenwood et al. (2018) (44), tDCS applied to the dorsal attention network (DAN) should facilitate externally directed cognition by enhancing DAN activity. In our study, however, tDCS reduced the presence of executive-control microstate D, suggesting a dampening rather than facilitation of externally oriented processing. By contrast, oscillatory protocols (tACS, otDCS) more closely aligned with Greenwood’s framework, suppressing DMN-related microstates (B, C) while enhancing MS D, thereby promoting a shift from internally focused to task-ready, externally oriented states.

Our findings also align with the proposed differential effects of tES on cognition (9), which suggest that distinct stimulation protocols may shape cognitive outcomes through distinct neurophysiological mechanisms. The consistent enhancement of MS D under oscillatory protocols (tACS, otDCS) points to a more direct modulatory route, whereby entrainment selectively strengthens executive-control networks. By contrast, tDCS effects appear to operate more indirectly, primarily via suppression of DMN-related microstates (B, C) and facilitation of MS A, reflecting enhanced sensory readiness and attention. The variability of tDCS effects often reported in the behavioral literature (45) may partly stem from processes leading to the suppression of MS D observed in our data, which could offset potential cognitive benefits by reducing executive-control engagement. In this light, oscillatory protocols may hold greater promise for producing reliable improvements in cognitive outcomes, as they more consistently foster task-ready, control-related network states.

Our results both converge with and extend prior tES–microstate findings. Earlier work has shown that tDCS can increase MS A (sensory/phonological) and MS D (cognitive control) while reducing MS C (self-referential), depending on stimulation site and population (21,22). In our study, we similarly observed facilitation of MS A but, in contrast, found suppression of MS D, an executive-control related state, suggesting that while tDCS reliably enhances early sensory-related states, its influence on higher-order control states is more variable. Oscillatory stimulation results were more consistent: γ-tACS has previously been shown to reduce MS C and enhance MS D, reflecting a shift from self-referential toward executive control networks (16,25). We extend this by demonstrating that tACS and otDCS suppression is not limited to MS C but also includes MS B, particularly under otDCS, indicating a stronger downregulation of visual-dominant states.

This resonates with findings from single-session rTMS studies, which similarly show that stimulation protocols, depending on stimulation site, exert distinct influences on microstates. Qiu et al. (2020) reported that 1 Hz rTMS over the motor cortex increased the duration of all microstates, consistent with a broad stabilization of brain states (46). In contrast, Croce et al. (2018) found that 1 Hz rTMS over the parietal cortex selectively reduced the occurrence of MS C, indicating targeted modulation of self-referential states, though earlier work also interpreted this effect as salience-related (14). Interestingly, this parietal effect parallels the reduction of MS C observed with γ-tACS over frontal–parietal regions (16,25) and with the effects of our theta-oscillatory protocols, suggesting that parietal stimulation, whether magnetic or electrical, consistently targets self-referential networks. In contrast, the tDCS pattern of effects in our study more closely resembles the general stabilization observed with motor cortex rTMS (8), underscoring the distinction between constant and oscillatory forms of stimulation.

The differential effects of constant and oscillatory stimulation on microstate dynamics can be interpreted in light of their distinct neurophysiological mechanisms. tDCS produces a tonic shift in neuronal membrane potential, biasing populations of neurons toward depolarization or hyperpolarization without imposing rhythmic structure (2,47,48). Such steady-state modulation is thought to stabilize baseline network configurations, which may explain the broad but relatively modest adjustments observed in our data. In contrast, tACS applies rhythmic alternating currents that interact with ongoing oscillatory activity, promoting phase alignment and frequency-specific entrainment of neural populations (48,49). This mechanism enables more selective reorganization of large-scale networks, consistent with the state-specific modulation of MS C and MS D we observed. The otDCS, combining a constant and an oscillatory component, likely exerts both tonic and phasic effects, which may account for its consistently strongest and most widespread influence on microstates. A recent report by Guo et al. (2025) further supports the view that otDCS combines the stabilizing effects of tDCS with the entrainment properties of tACS (8). Authors directly compared 40 Hz otDCS, tACS, and tDCS over the motor cortex and showed that otDCS produced the strongest and most sustained increases in corticospinal excitability, as well as the most pronounced enhancement of large-scale network connectivity measured with TMS–EEG. These effects were interpreted as reflecting a synergistic mechanism, whereby otDCS simultaneously induces tonic membrane polarization and oscillatory entrainment, thereby amplifying NMDA receptor–dependent plasticity. Our findings converge with this interpretation: otDCS consistently exerted the strongest influence on EEG microstates, modulating all canonical states and producing both tDCS-like stabilization of sensory-related activity and tACS-like reorganization of self-referential and executive-control networks. Supporting evidence from animal models also shows that otDCS, when paired with tDCS, more powerfully enhances endogenous oscillatory activity compared to tDCS alone, especially when matched to intrinsic frequency bands (50). This indirectly supports our finding that otDCS exerts the most robust modulation of EEG microstates. These distinctions suggest that microstate analysis is sensitive not only to stimulation site and frequency but also to the underlying physics of how different current waveforms shape cortical network dynamics.

Recent evidence from high-definition transcranial direct current stimulation (HD-tDCS) further supports our interpretation. Zhou et al. (2025) found that anodal HD-tDCS over the right dorsolateral prefrontal cortex (dlPFC) reduced MS B and enhanced MS D, mirroring our findings under oscillatory protocols (51). This indicates that the site and focality of stimulation can influence the effects of constant current protocols. While traditional tDCS induces broad shifts, HD-tDCS may selectively target prefrontal networks, modulating oscillatory dynamics and enhancing executive functions. Overall, waveform, frequency, and focality work together to shape network reorganization, with microstates serving as a sensitive measure of these effects. Notably, we delivered both tACS and otDCS at a theta frequency that was individual to each participant. This frequency-personalization approach is grounded in the idea that stimulation effects would be maximized when the external current matches intrinsic oscillatory dynamics, thereby increasing the likelihood of phase alignment and effective entrainment (52–55). Importantly, we targeted the theta range because of its central role in large-scale brain communication, particularly in coordinating memory and executive-control processes through long-range frontoparietal and hippocampal–cortical networks (56,57). Unlike alpha, which primarily supports sensory inhibition, or gamma, which is more closely linked to local synchronization, theta oscillations provide a temporal scaffold for long-range integration across distributed networks (58–60). Thus, delivering stimulation at the ITF ensured both frequency specificity and individualized targeting, which may explain why ITF-oscillatory protocols showed selective and consistent modulation of self-referential (MS C) and executive-control (MS D) microstates, in contrast to the more diffuse effects of tDCS. Namely, aligning stimulation with oscillations endogenous to the targeted network likely amplifies resonance effects and facilitates more efficient large-scale reorganization, consistent with evidence that network entrainment is stronger when stimulation frequency is individually tuned (61).

The observed stimulation-induced modulations also have broader functional and clinical significance. Facilitation of MS A, reflecting sensory readiness and arousal, may help explain why tDCS protocols often produce modest but reliable effects on pain perception and basic sensory processing (21). Suppression of MS B and MS C by oscillatory stimulation suggests a reduction of self-focused and autobiographical mentalization, processes often exaggerated in conditions such as depression and rumination (13,42). Conversely, the enhancement of MS D under tACS and otDCS indicates a strengthening of executive and attentional control networks, consistent with findings that γ-tACS improves working memory performance (16). Taken together, these results highlight that oscillatory stimulation may be particularly suited for clinical contexts where self-referential dominance and executive deficits coexist, such as in depression, schizophrenia, or OCD, whereas constant currents may be more effective in modulating sensory readiness or stabilizing overall network dynamics.

Overall, this study makes several broader contributions. Mechanistically, it provides the first direct evidence that constant and oscillatory tES protocols differentially reorganize large-scale brain networks, with otDCS bridging polarity-driven and oscillatory mechanisms. Methodologically, it establishes EEG microstate analysis as a sensitive, temporally precise tool for detecting network-level reorganization induced by neuromodulation. Unlike EEG analyses focused on oscillatory power or connectivity, which capture frequency-specific or pairwise interactions, microstates reflect the rapid succession of whole-brain configurations and thus provide a complementary window into the global coordination of neural activity, making it particularly suited for capturing the broad and distributed effects expected from brain stimulation. Translationally, the results offer proof of concept that microstates from resting state EEG can serve as practical biomarkers for tracking and differentiating the large-scale effects of distinct tES modalities.

### Limitations and Future Directions

Several limitations should be acknowledged. First, our analysis was restricted to the four canonical microstates (A–D). This means that potential distinctions between MS C (self-referential mentation) and MS E (salience-related activity) could not be examined (13), and some of the effects attributed to MS C may partly reflect overlapping salience-related dynamics. To our knowledge, no existing brain stimulation studies have reported effects on MS E, as most have similarly adopted the four-class solution. Second, the study was conducted in healthy young adults, which limits generalizability to clinical or aging populations where stimulation effects may differ. Third, only offline resting-state effects were assessed; the extent to which these modulations translate into task-related brain dynamics or behavioral outcomes remains an open question. Our results suggest that stimulation at individual theta frequencies may enhance the consistency and potency of oscillatory tES effects on large-scale brain networks, but this interpretation should be considered with caution, given that we did not directly compare individualized with fixed-frequency protocols. Finally, otDCS is a relatively new protocol, and its mechanisms of action require further validation. Future work should therefore examine microstates using higher-class solutions, extend investigations to clinical groups, and combine EEG with multimodal imaging and behavioral assessments to more fully capture the functional and therapeutic relevance of stimulation-induced network changes.

## Conclusion

This study demonstrates that EEG microstates provide a sensitive framework for disentangling the network-level effects of different tES modalities. While tDCS primarily modulated sensory-related states in a manner consistent with network stabilization, oscillatory protocols (particularly otDCS) more strongly reorganized self-referential and cognitive control-related states. By conducting the first within-subject, multimodal comparison of tDCS, tACS, and otDCS, we identified both shared and modality-specific influences on canonical microstates, with otDCS exerting the most robust and widespread effects. These findings advance mechanistic understanding of how distinct current waveforms shape large-scale brain dynamics and establish EEG microstates as a promising biomarker for differentiating, optimizing, and clinically monitoring neuromodulation strategies.

## Supplementary material

**Table S1.**
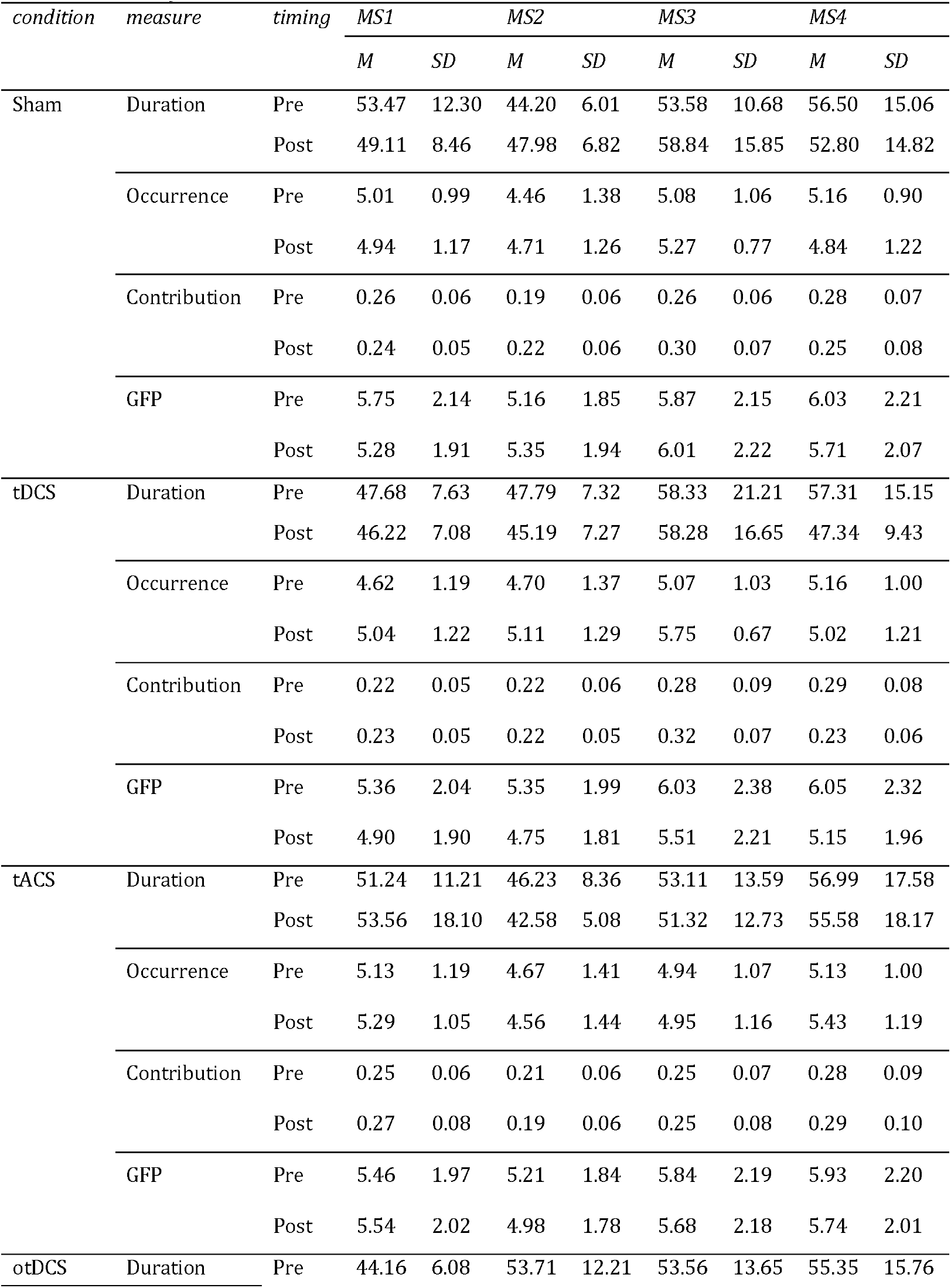
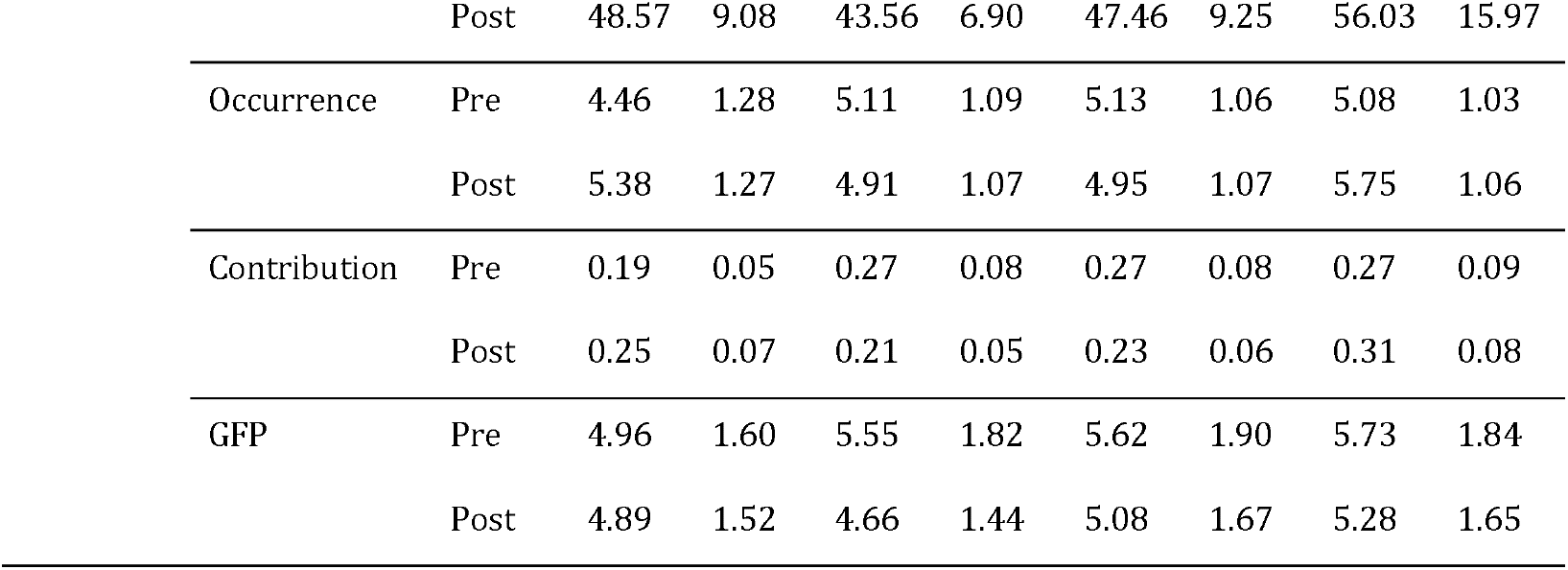
Descriptive statistics for microstates measures pre- and post-stimulation across conditions (Sham, tDCS, otDCS, tACS)

**Table S2.** Statistical outcomes of tES effects on microstates.

| *Condition* | *MS* | *Measure* | *condition* |  |  | *time* |  |  | *condition x time* |  |  |
| --- | --- | --- | --- | --- | --- | --- | --- | --- | --- | --- | --- |
| | | | *F*<sub>(1,123)</sub> | *p* | $\eta^2$ | *F*<sub>(1,123)</sub> | *p* | $\eta^2$ | *F*<sub>(1,123)</sub> | *p* | $\eta^2$ |
| tDCS | MS1 | Duration | **18.58** | **<.001** | **.13** | **8.34** | **.005** | **.06** | 2.07 | .152 | .02 |
|  |  | Occurrence | 1.30 | .257 | .01 | 2.07 | .153 | .02 | **4.05** | **.046** | **.03** |
|  |  | Contribution | **13.34** | **<.001** | **.10** | 0.83 | .365 | .01 | **6.07** | **.015** | **.05** |
|  |  | GFP | **10.58** | **.001** | **.08** | **15.51** | **<.001** | **.11** | 0.00 | .959 | .00 |
|  | MS2 | Duration | 0.38 | .537 | .00 | 0.83 | .363 | .01 | **24.53** | **<.001** | **.17** |
|  |  | Occurrence | **8.25** | **.005** | **.06** | **8.53** | **.004** | **.06** | 0.55 | .461 | .00 |
|  |  | Contribution | **7.30** | **.008** | **.06** | **10.29** | **.002** | **.08** | **4.18** | **.043** | **.03** |
|  |  | GFP | *3.61* | *.060* | *.03* | **3.97** | **.049** | **.03** | **14.62** | **<.001** | **.11** |
|  | MS3 | Duration | 1.30 | .257 | .01 | 2.00 | .160 | .02 | 2.08 | .152 | .02 |
|  |  | Occurrence | **5.30** | **.023** | **.04** | **18.74** | **<.001** | **.13** | **5.84** | **.017** | **.05** |
|  |  | Contribution | 3.23 | *.075* | *.03* | **11.82** | **<.001** | **.09** | 0.03 | .856 | .00 |
|  |  | GFP | 1.63 | .204 | .01 | 2.10 | .150 | .02 | **6.48** | **.012** | **.05** |
|  | MS4 | Duration | 2.93 | *.089* | *.02* | **25.24** | **<.001** | **.17** | **5.32** | **.023** | **.04** |
|  |  | Occurrence | 0.55 | .458 | .00 | *3.88* | *.051* | *.03* | 0.60 | .440 | .00 |
|  |  | Contribution | 0.33 | .568 | .00 | **27.43** | **<.001** | **.18** | 1.28 | .261 | .01 |
|  |  | GFP | **4.73** | **.032** | **.04** | **23.63** | **<.001** | **.16** | **5.21** | **.024** | **.04** |
| otDCS | MS1 | Duration | **21.55** | **<.001** | **.15** | 0.00 | .981 | .00 | **17.11** | **<.001** | **.12** |
|  |  | Occurrence | 0.21 | .649 | .00 | **12.40** | **.001** | **.09** | **16.79** | **<.001** | **.12** |
|  |  | Contribution | **9.53** | **.003** | **.07** | 5.55 | **.020** | **.04** | **31.25** | **<.001** | **.20** |
|  |  | GFP | **22.20** | **<.001** | **.15** | **4.56** | **.035** | **.04** | 2.62 | .108 | .02 |
|  | MS2 | Duration | **6.53** | **.012** | **.05** | **10.24** | **.002** | **.08** | **49.04** | **<.001** | **.29** |
|  |  | Occurrence | **12.99** | **<.001** | **.10** | 0.04 | .850 | .00 | *3.58* | *.061* | *.03* |
|  |  | Contribution | **15.20** | **<.001** | **.11** | 3.76 | *.055* | *.03* | **28.53** | **<.001** | **.19** |
|  |  | GFP | 1.37 | .244 | .01 | **8.35** | **.005** | **.06** | **19.87** | **<.001** | **.14** |
|  | MS3 | Duration | **15.36** | **<.001** | **.11** | 0.08 | .772 | .00 | **15.26** | **<.001** | **.11** |
|  |  | Occurrence | 1.45 | .230 | .01 | 0.00 | .960 | .00 | *2.77* | *.099* | *.02* |
|  |  | Contribution | **11.00** | **.001** | **.08** | 0.02 | .877 | .00 | **12.08** | **<.001** | **.09** |
|  |  | GFP | **16.31** | **<.001** | **.12** | 1.88 | .173 | .02 | **5.62** | **.019** | **.04** |
|  | MS4 | Duration | 0.74 | .393 | .00 | 1.54 | .217 | .01 | *3.24* | *.074* | *.03* |
|  |  | Occurrence | **11.66** | **.001** | **.09** | 2.13 | .147 | .02 | **16.54** | **<.001** | **.12** |
|  |  | Contribution | **9.53** | **.002** | **.07** | 0.02 | .881 | .00 | **16.52** | **<.001** | **.12** |
|  |  | GFP | **7.18** | **.008** | **.06** | **8.00** | **.005** | **.06** | 0.20 | .652 | .00 |
| tACS | MS1 | Duration | 0.63 | .427 | .01 | 0.53 | .466 | .00 | **5.77** | **.018** | **.04** |
|  |  | Occurrence | **4.95** | **.028** | **.04** | 0.15 | .695 | .00 | 1.08 | .300 | .01 |
|  |  | Contribution | *3.06* | *.083* | *.02* | 0.19 | .663 | .00 | **5.86** | **.017** | **.05** |
|  |  | GFP | 0.01 | .919 | .00 | *3.52* | *.063* | *.03* | **7.03** | **.010** | **.05** |
|  | MS2 | Duration | **5.15** | **.025** | **.04** | 0.01 | .924 | .00 | **25.00** | **<.001** | **.17** |
|  |  | Occurrence | 0.07 | .788 | .00 | 0.40 | .526 | .00 | 2.70 | .103 | .02 |
|  |  | Contribution | 2.19 | .141 | .02 | 0.55 | .461 | .00 | **19.94** | **<.001** | **.14** |
|  |  | GFP | 2.74 | .100 | .02 | 0.04 | .840 | .00 | **5.01** | **.027** | **.04** |
|  | MS3 | Duration | **8.39** | **.004** | **.06** | 1.57 | .212 | .01 | **6.52** | **.012** | **.05** |
|  |  | Occurrence | **4.63** | **.033** | **.04** | 0.81 | .369 | .01 | 0.69 | .407 | .01 |
|  |  | Contribution | **10.24** | **.002** | **.08** | 1.94 | .166 | .02 | **4.48** | **.036** | **.04** |
|  |  | GFP | 2.35 | .128 | .02 | 0.01 | .933 | .00 | 1.82 | .179 | .01 |
|  | MS4 | Duration | 1.03 | .313 | .01 | 2.52 | .115 | .02 | 0.50 | .481 | .00 |
|  |  | Occurrence | **4.77** | **.031** | **.04** | 0.01 | .945 | .00 | **6.04** | **.015** | **.05** |
|  |  | Contribution | **4.89** | **.029** | **.04** | 1.68 | .198 | .01 | **4.70** | **.032** | **.04** |
|  |  | GFP | 0.11 | .743 | .00 | **5.32** | **.023** | **.04** | 0.38 | .537 | .00 |

*Note.* $p < .05$ – bolded, $p < .10$ – italicized

**Figure S1.**
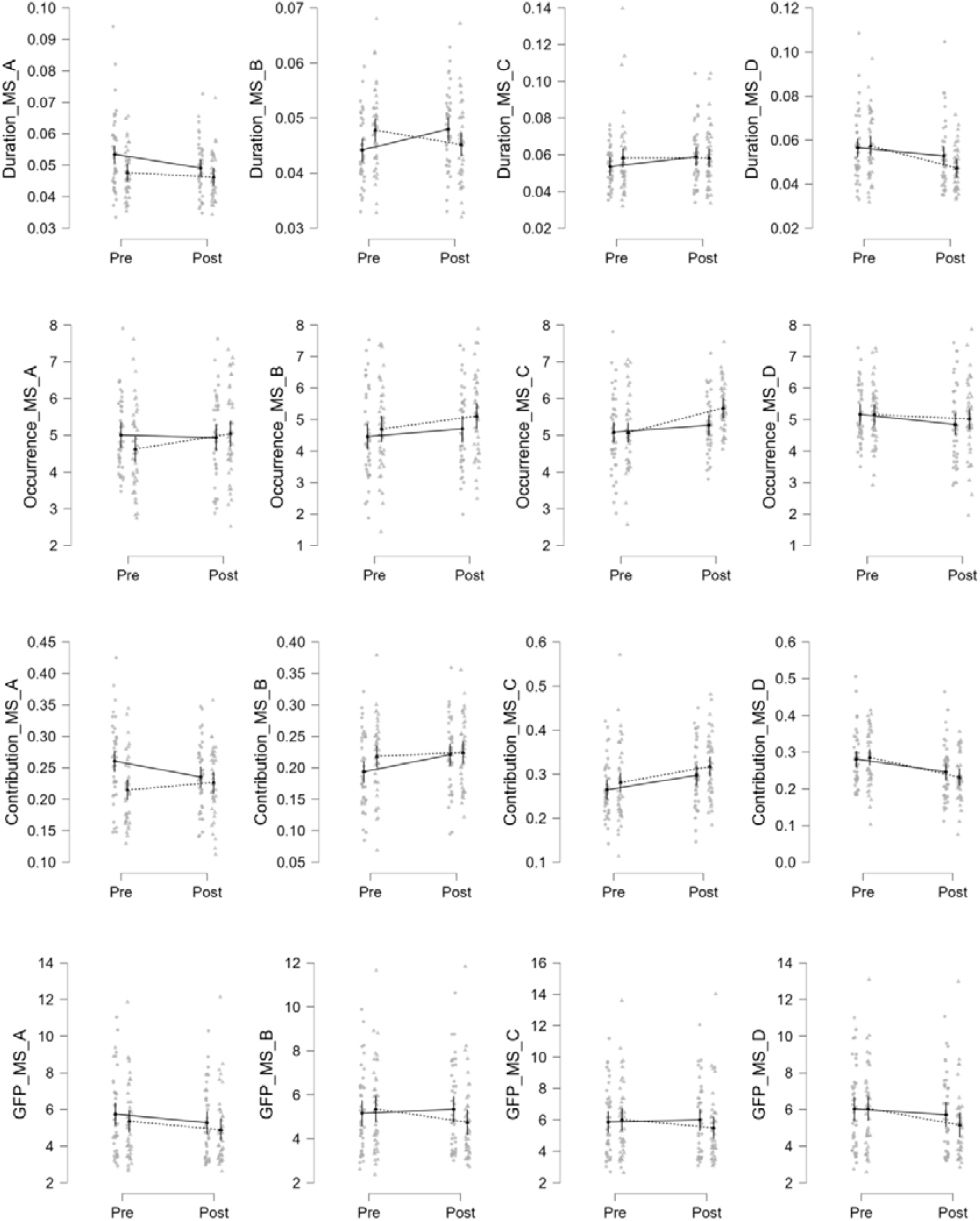
The effects of **tDCS** and time of assessment on MS measures.

**Figure S2.**
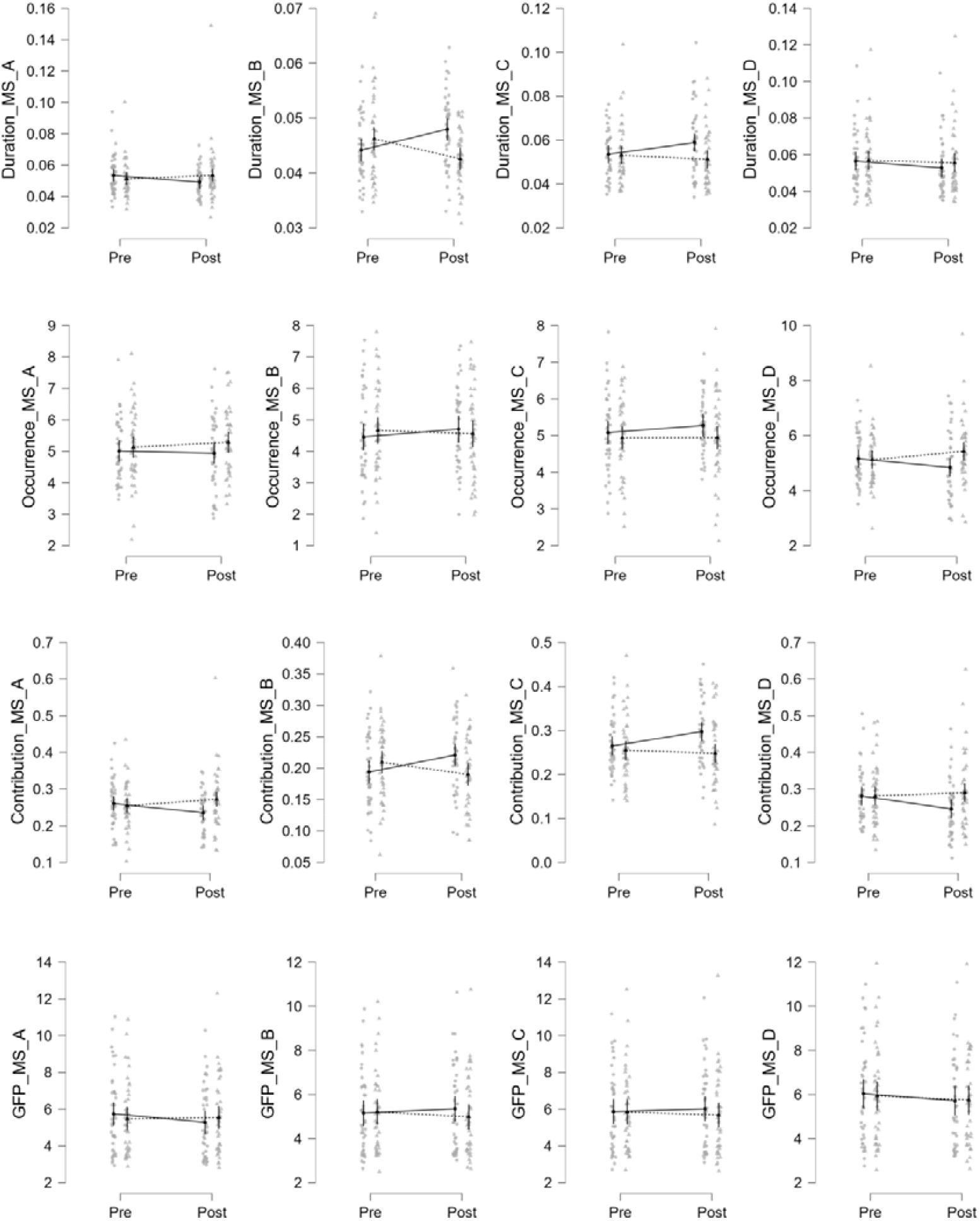
The effects of **tACS** and time of assessment on MS measures.

**Figure S3.**
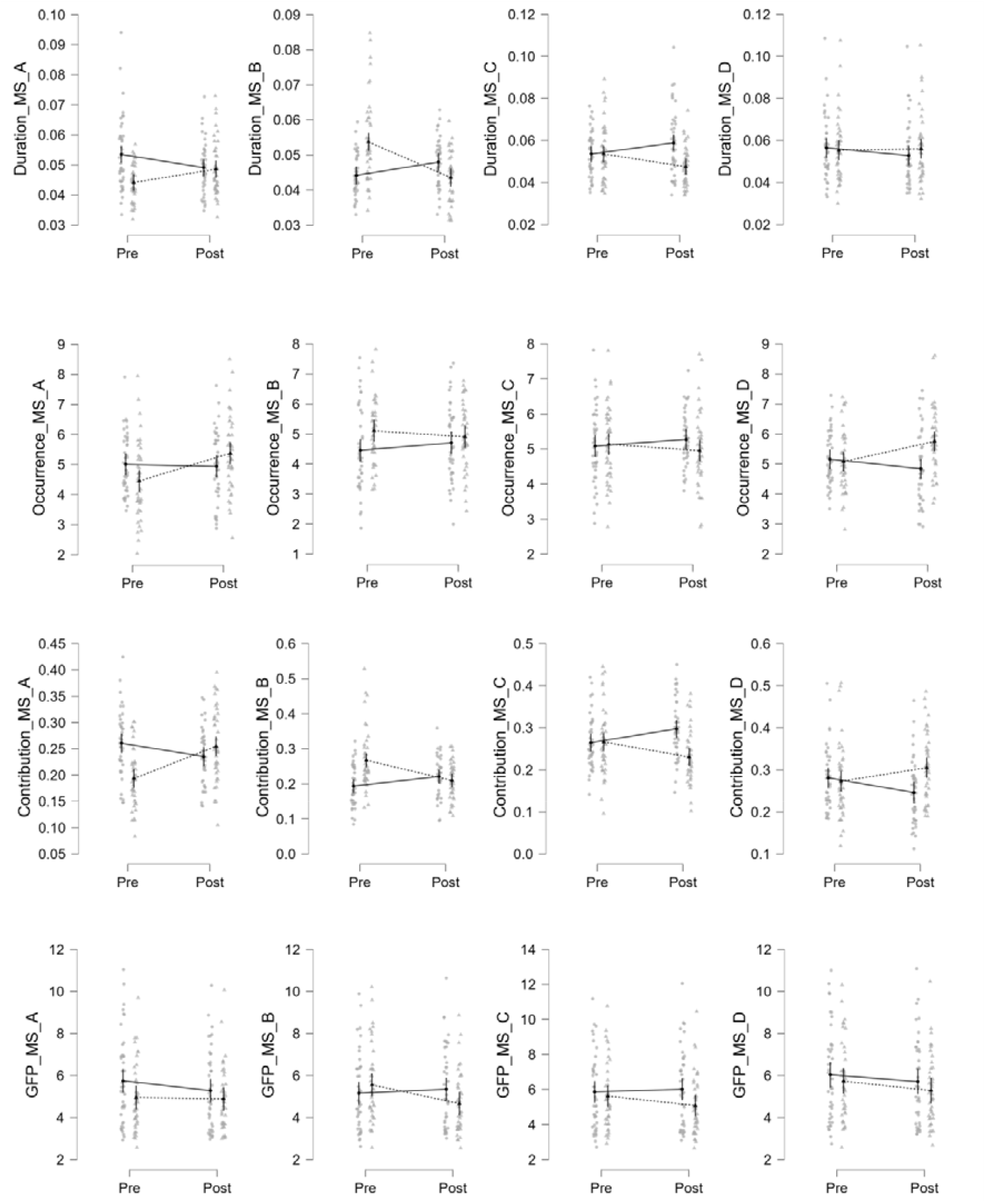
The effects of **otDCS** and time of assessment on MS measures.

## Notes

<u>Funding:</u> The data for this study have been collected under the MEMORYST grant awared by Science Fund of Republic of Serbia. MŽ and JB receive institutional support from the Ministry of Science, Technological Development and Innovation of the Republic of Serbia (451-03-137/2025-03/200163; 451-03-136/2025-03/200015). IGB, MŽ, SF and JB are engaged in EU-funded HORIZON Collaboration and Support Action TWINNIBS (101059369).

### Competing Interest Statement

The authors have declared no competing interest.

